# Protein Engineering with A Glycosylation Circuit Enables Improved Enzyme Characteristics

**DOI:** 10.1101/2021.11.15.468597

**Authors:** Eray Ulaş Bozkurt, İrem Niran Çağıl, Ebru Şahin Kehribar, Musa Efe Işılak, Urartu Özgür Şafak Şeker

## Abstract

Protein glycosylation is one of the most crucial and common post-translational modifications. It plays a fate-determining role and can alter many properties of proteins, making it an interesting for many biotechnology applications. The discovery of bacterial glycosylation mechanisms, opened a new perspective and transfer of *C*.*jejuni* N-linked glycosylation into laboratory work-horse *E. coli* increased research pace in the field exponentially. It has been previously showed that utilizing *N-Linked Glycosylation*, certain recombinant proteins have been furnished with improved features, such as stability and solubility. In this study, we utilized *N-linked Glycosylation* to glycosylate alkaline phosphatase (ALP) enzyme in *E. coli* and investigate the effects of glycosylation on an enzyme. Considering the glycosylation mechanism is highly dependent on the acceptor protein, ALP constructs carrying glycosylation tag at different locations of the gene has been created and glycosylation rates have been calculated. The most glycosylated construct has been selected for comparison with the native enzyme. We investigated the performance of glycosylated ALP in terms of its thermostability, proteolytic stability, tolerance to suboptimal pH and under denaturing conditions. Studies showed that glycosylated ALP performed remarkably better at optimal and harsh conditions Therefore, *N-linked Glycosylation* mechanism can be employed for enzyme engineering purposes and is a useful tool for industrial applications that require enzymatic activity.

## Introduction

Protein glycosylation, which refers to attachment of sugar residues to proteins, is one of the most prevalent, diverse and crucial post-translational modifications (Kightlinger et al., 2020). The phenomenon is found in all domains of life(Dell et al., 2010) and over 50% of eukaryotic proteome is known to be glycosylated.(Baker et al., 2013). Glycosylation plays a significant roles and it can alter many features of proteins, including stability and activity, therefore atracts wide attention from scientific community (Elliott et al., 2003; Li et al., 2006). Although, recombinant proteins are frequently utilized in medical and industrial applications (Wurm, 2019), many proteins often suffer from low stability under suboptimal thermal and chemical conditions and exhibit low solubility. They also may aggregate over time which may cause decreased efficiency and increased immunogenicity. Such factors limit the efficient usage of recombinant proteins in many applications and manipulation of these properties could create new opportunities in medical and industrial approaches (Ma et al., 2020).

Protein glycosylation, once thought to be unique for eukaryotes, was later discovered in Bacteria and Archaea (Abu-Qarn et al., 2008). Although glycosylation systems are more diverse in eukaryotes, N-linked and O-linked glycosylation are commonly found in bacteria (Nothaft and Szymanski, 2010). N-linked glycosylation refers to addition of glycans to the nitrogen group of asparagine residues, whereas O-linked glycosylation is the addition to the hydorxyl oxygen on serine or threonine residues (Nothaft and Szymanski, 2010). N-linked glycosylation first described in *Campylobacter jejuni* (*C. Jejuni)* (Szymanski et al., 1999) and shown that *C. Jejuni* utilizes pgl (protein glycosylation) mechanism to attach glycan groups *en bloc* to more than 65 proteins in the periplasm (Scott et al., 2011). The glycosylated proteins play role on contributing to the fitness of the bacterium in the gut and protecting it from proteases (Alemka et al., 2013; Lu et al., 2015). This discovery speed up the progress being made in the field and many other findings followed (Baker et al., 2013). Today, it is known that N-linked glycosylation at least 49 species possess the compounds of pgl pathway (Wacker et al., 2002). N-linked glycsoylation can also occur as a sequential transfer rather than *en bloc*. Such glycosylation machinery was discovered in *Haemophilus influenzae* (*H. influenzae). H. influenzae* utilizes this mechanism to glycosylate two of its proteins, HMW1 and HMW2 which are secreted to outer membrane and contributes to adhesiveness to the host surfaces (St Geme et al., 1993). Unlike many bacteria, *Actinobacillus pleurop-neımoniae* possess soluble enzymes and is able to glycosylate proteins in the cytoplasm(Schwarz et al., 2011). In addition to N-linked glycosylation, O-linked glycosylation is also reported in certain bacteria, including *Neisseria meningitidis* and *Neisseria gonorrhoeae*. The process is similar to N-linked glycosylation and occurs in the cytoplasm. Despite the efforts to enlighten novel pathways from different strains, *C. jejuni*’s N-linked glycosylation remains to be most characterized pathway (Baker et al., 2013).

C. jejuni N-linked glycosylation mechanism is composed of 13 genes encoding enzymes for synthesis of glycan and *en bloc* transfer of the produced glycan moiety onto acceptor protein. N-glycan synthesis is initiated by formation of uridine diphosphate (UDP)-activated-*N*-acetylglucosamine (UDP-GlcNAc) on the cytoplasmic side of the cell. PglF enzyme, a C6 dehydratase, converts UDP-GlcNAc to UDP-2-acetamido-2,6-dideoxy-a-D-xylo-4-hexulose, which is followed by formation of UDP-4-amino-4,6-dideoxy-α-D-GlcNAc by PglE. Then, PglD transfers the acetyl group. PglC links UDP-diNAcBac to a lipid-linked precursor, undecaprenyl phosphate (Und-P). PglA and PglJ catalyzes sequential reactions to add α 1,3- and α1,4-linked N-acetylgalactosamine. Then, PglH adds three α1,4-linked GalNAc. In the final step, PglI attaches β1,3-linked glucose and completes the synthesis. The lipid linked oligosaccharide (LLO) is now flipped from the cytoplasmic side to periplasmic side of the cell by PglK, an ABC transporter enzyme. Then, PglB, the oligosaccharyltransferase, transfers the heptasaccharide onto the glycosylation sequon (D/E-X_1_-N-X_2_-(S/T) where X_1_ and X_2_ cannot be proline)(Tan et al., 2015) The discovery of the glycosylation machinery of *Campylobacter jejuni (C. Jejuni)* (Young et al., 2002) and its recapitulation in laboratory workhorse *E. coli* has greatly leveraged the possibilities in glycoengineering (Wacker et al., 2002).

Glycoengineering approach has been widely used in many different organisms to develop better therapeutics. In mammalian cells, one of the major aims is to obtain more homogeneous products in terms of N-linked glycosylation patterns since heterogeneity adds complexity during optimization (Rich and Withers, 2009). Researchers have adopted several approaches such as zinc finger nucleases and CRISPR-Cas9 to knock-out glycosyltrannsferase enzyme to reduce glycan diversity on final product (Pereira et al., 2018; Tian et al., 2019; Yang et al., 2015). Glycoengineering was also employed to develop vaccines and glycomaterials utiling different host models.(Kightlinger et al., 2020). One example is the utilization of glycoengineering to produce conjugate vaccines. Conjugate vaccines are produced by covalently linking adjuvant to a carrier protein. Utilization of glycosylation offers a easy and cost-effective alternative to vaccine production procedure. Recombinant glycosylation can also be used to alter biophysical and pharmokinetic properties. One study showed that addition of glycosylation sites leads to increased circulation times and binding abilities (Elliott et al., 2003). Another study reported that glycosylation of a single-chain antibody improved the solubility and proteolytic stability. (Lizak et al., 2011). Lastly, to produce glycomaterials for enhanced properties, biofilm proteins were N-linked glycosylated. In the study, it has been shown that addition of N-glycans to biofilm proteins lead to a protein-based biomaterial with enhanced adsorption properties (Sahin Kehribar et al., 2021). Overall, glycoengineering can contribute to many fields such as vaccine development, therapeutics as well as industrial processes and can serve as a cost-effective and efficient alternative to current methods.

In this study, we employed N-linked glycosylation mechanism of *C. jejuni* to recombinantly produce glycosylated enzymes in *E. coli*. As the proof of concept, we chose alkaline phosphatase enzyme (ALP) and aim to investigate the effects of N-linked glycosylation on the behavior of the enzyme. We also examine the secondary structure analysis under different conditions and assessed the enzymatic activity differences upon glycosylation.

## Materials & Methods

### Plasmid Construction and Bacterial Growth Conditions

Plasmids and primers were designed using Benchling. PCR reactions were performed using Q5 High-Fidelity DNA Polymerase (New England Biolabs Inc.) by following the manufacturer’s instructions. Primers that were used in PCR reactions were ordered from Oligomer. Annealing temperatures for PCR reactions were calculated using the NEB Tm Calculator tool.

*phoA*-DQNAT pET22b plasmid was digested with BamHI to insert *pglB* gene. Amplification and insertion of *pglB* gene encoding for pglB enzyme in the pgl pathway was performed via Gibson Assembly. The gene was amplified using the primers: 5’CTAGAATTAAAGAGGAGAAAGGTACGCGGATCCATGTTGAAAAAAGAGTATTT AAAAA3’ and 5’CCAGTGCAATAGTGCTTTGTTTCATGTTTCTCCTCTTTAATACTAGTTTAAATTTTAAGTTTAAAAACCTTAG3’.

PCR products were mixed with 6X Purple Loading Dye (New Engliand Biolabs Inc., B7024S) and run in 1% agarose gel. 1kb+ Ladder (New England Biolabs Inc., N3200L) was utilized to track DNA length. To prepare agarose gels, 0.6g of agarose was dissolved in 60 mL 1X TAE buffer. Electrophoresis was performed for 30 min at 140V. Bands confirmed by electrophoresis was extracted using Macherey-Nagel GmbH & Co. gel extraction and PCR clean-up kit, following manufacturer’s instructions and yield was measured using Nanodrop 2000 spectrophotometer (Thermo Fisher, ND2000).

In restriction digestion reactions New England Biolab enzymes were used and manufacturer instructions were followed. Fragments were run on agarose and extracted as mentioned above. Cloning products were transformed into chemically competent cells. To perform transformation, chemically competent cells were thawed on ice. Then, cloning products were added onto cells and incubated for 20 minutes on ice. Heat-shock was applied at 42°C for 30 seconds and transferred to ice again for 2 minutes. 1 mL of LB medium was added and cells were incubated at 37°C for 1 hour. After incubation, centrifugation was performed at 8000 G for 5 minutes and supernatant was discarded. Pellet was resuspended in residual supernatant and laid on agar plates with appropriate antibiotics. Agar plates were incubated at 37°C O/N. After incubation, colonies were screened with colony PCR. Positive colonies were inoculated into LB medium with appropriate antibiotics. Plasmids were isolated using Gene-JET Miniprep Kit (Thermo Scientific, K0503). Isolated plasmids were verified with Sanger sequencing.

### SDS-PAGE, Coomassie Blue Staining, and Immunoblotting

SDS-PAGE was performed both to verify purity of the proteins and to observe the effects of proteolytic cleavage. For the verification part, 20 μl of 2 μg/ml protein in 25 mM Tris-HCl were mixed with 6X SDS Loading Dye (375 mM Tris-HCl (pH 6.8), 9% (w/v) SDS, 50% (v/v) glycerol, 0.03% (v/v) bromophenol) and denatured at 95°C for 5 min. SDS gel was prepared by BioRad SDS Gel Casting System and 20 μl of the mixture was loaded onto the gel. The proteins were run on the gel for nearly 2 hours by applying 120-150 V. For observation of effects of the proteolytic cleavage 20 μl of 100 μg/ml protein in Proteinase K, 20 μl of the purified protein in urea, and 20 μl of the purified protein in sodium phosphate buffer were mixed with 6X SDS Loading Dye and same procedures were applied for the electrophoresis part. Coomassie Blue Staining was performed, and the gel was visualized by Image Lab Software (Bio-Rad).

Whole-cell western and lectin blots were applied to different concentrations of recombinant protein producing cells to verify the presence of recombinant proteins and their glycosylated forms by applying the same procedures. The proteins then transferred into polyvinylidene difluoride (PVDF) membrane (Thermo Fisher Scientific 88518) after running. PVDF membrane was activated by methanol and then put in Turbo Transfer Buffer (Bio-Rad). Transblot Turbo Transfer System (Bio-Rad) was used for semi-dry transferring in 7 min.

For Western Blotting, 1x Tris Buffer Saline with 0.1% of Tween (TBS-T) containing 5% skimmed milk was used as blocking solution. The membrane was blocked with the blocking solutions for 2 hours at room temperature. Primary antibody (Anti-His Mouse PTGLAB 66005) was diluted at 1:10000 in blocking solution and the incubation step was done for 1 hour at room temperature. The membrane was washed with 1x TBS-T (0.1%) three times for 5-, 10-, and 10-min. Secondary antibody (HRP conjugated goat anti-mouse) (Abcam ab6789-1 MG) incubation was performed for 2 hours at room temperature by diluting it at 1:10000 in blocking solution. Finally, the wash steps were repeated. Visualization of the proteins on the membrane was performed by Enhanced chemiluminescence (ECL) (Bio-Rad 170-5060) via Image Lab Software (Bio-Rad).

For Lectin Blotting, blocking solution was filtered 1x TBS with 0.05% of Tween (TBS-T) containing 3% BSA. The membrane was blocked with the blocking solution for 1 hour at room temperature. Primary antibody was diluted at 1:5000 in blocking solution and its incubation was done for 2 hours at room temperature. Wash steps were performed by 1xTBS-T (0.05%) five times for 5 min each. The visualization steps were performed also by ECL.

### Recombinant Protein Expression and Extraction

Cells containing plasmids were inoculated from the stock in ZMY052 autoinduction media. After a 20-24 hour incubation, protein extraction steps were performed. Isolated cells by centrifuging at 3500 G for nearly 30 min were resuspended with 10 mM imidazole buffer (20 mM Sodium Phosphate, 0.5 M NaCl, 10 mM imidazole, pH 7.4). Proteases were inhibited with the addition of 1 mM phenylmethanesulfonylfluoride (PMSF) (AMRESCO Inc.) and cells were lysed by sonication. In the sonication step, 30 sec pulse-on and 59 sec pulse-off with amplitude 0.35 were applied 5 times. To collect recombinant proteins, lysed cells were centrifuged at 12000 rpm for 1 hour and total protein was extracted as the supernatant and stored at +4°C.

### Recombinant Protein Purification by Cobalt Resin

For the purification of recombinant proteins, 10 mM imidazole was used as binding buffer. The lysis supernatant containing total protein was added on to the resin and the mixture was rotated for 1 h to provide the binding of His-Tagged ALP to the resin. Unbound proteins were discarded and the resin was washed three times with 10 mM imidazole. Bound proteins were eluted three times with 150 mM imidazole (20 mM Sodium Phosphate, 0.5 M NaCl). Buffers in which proteins found were exchanged with 25 mM Tris-HCl (pH 8), Sodium Phosphate Buffer (1 M Na2HPO4, 1 M NaH2PO4), 8 M Urea (Sigma Aldrich, 51457), and Guanidine Hydrochloride (Serva, 24205.02) via Hi-Trap Desalting column (Sigma Aldrich GE17140801) for further experiments. Quantification of the recombinant proteins was performed by Pierce™ BCA Assay (Thermo Fisher Scientific 23225) according to manufacturer’s instructions.

### Glycosylation Rate Estimations

The glycosylation rate of the constructs was determined using Western Blotting analysis. The analysis of bands was conducted using Vilber Evolution Capt Edge software after viewing with the Vilber Lourmat FUSION SOLO 6 imaging equipment. In the software, quantification application was selected. Background signals were substracted from the image and equal area was selected for most precise calculation.

### ALP Activity Assay

Enzymatic activity was determined for both the wild type and the glycosylated form of ALP by measuring the catalysis of pNPP substrate. pNPP reaction buffer (0.1 M Glycine, 1 mM MgCl2, 1 mM ZnCl2) was used to make pNPP substrate solutions. The Michaelis-Menten curve was created using various pNPP concentrations (0 mM, 1 mM, 2 mM, 3 mM, 4 mM, and 5 mM). The stability and activity variations of these enzymes were determined by calculating and comparing their enzymatic activity in various conditions. For this purpose, protein activities incubated at 37°C for 10 min and at 55, 75, and 95°C for 15, 30, 60, 120 min were measured to observe the effects of increasing temperature and incubation times on them. Incubations were performed on a thermal cycler (Bio-Rad). 50 ul of proteins diluted at 2 ug/ml with 25 mM Tris-HCl (pH 8) were mixed with 50 ul of pNPP substrates in this part of the experiment.

Effects of pH on enzymatic activities of recombinant proteins were calculated by measuring the catalysis of pNPP substrate. Therefore, proteins were diluted to 2 ug/ml with 25 mM Tris-HCl having pH 5, 6, 7, 8, 9, 10, 11. In this stage of the experiment, 50 ul of proteins were combined with 50 ul of pNPP substrates.

Treatments with proteinase K and urea were applied to test the stabilities of the enzymes. Thus, proteinase K concentration (100 uM, 20 uM, 10 uM, 5 uM, 0.5 uM, 0.05 uM, 0.005 uM, 0.0005 uM) and incubation time variants (60, 120, 240 min) were applied. After incubation at 37°C, 5 ul of Proteinase K were added to 50 ul of 2 ug/ml protein. Additionally, activities of 50 ul of 2ug/ml protein were measured in different urea concentrations (8M, 1M, 0.1M, 0.01M, 0.001M) after an incubation process lasting 1 hour at 37°C.

The reaction rate was estimated for all of the conditions mentioned above using the pNP standard curve and the absorbance of the well at 405 nm was measured at 37°C.

### Circular Dichroism

CD (JASCO J-815) studies were carried out to observe the changes in secondary structures of recombinant proteins with temperature, pH, and the presence of proteinase or denaturant. 1 uM in Tris buffer (pH 8.0) of each proteins were used for the experiments except pH changes. For pH, sodium phosphate buffer was used. Circular dichroism, voltage, and absorbance were chosen as the three channels. The wavelength was set to 190-250 nm, with 4 sec Digital Integration Time (D.I.T.), 1 nm band width, wavelength range, standard sensitivity, 100 nm/min scanning speed, 3 repeats accumulation mode.

## Results

### Expression of ALP and Glycosylated ALP in *E. Coli*

In this study, we focused on the effect of *N-linked glycosylation* of *C. Jejuni* on alkaline phosphatase enzyme’s behaviour. Although ALP is widely studied for decades, recombinant glycosylation of an enzyme and the effect of glycosylation has not been investigated before. By making an analogy to eukaryotic glycosylation, it is anticipated that recombinantly glycosylated enzymes will perform better at certain conditions, compared to the native enzyme (Figure 1A). Given that success of glycosylation event depends on the accessiblity of the recognition motif by PglB, two constructs were designed that carries glycosylation motif at N-terminus (DQNAT-ALP) and C-terminus (ALP-DQNAT). Also, another construct with two recognition motifs at C-terminus (ALP-DQNAT(x2)) was also designed. Designed constructs were co-transformed into cells with pgl-pACYC plasmid as well as without pgl-pACYC as a control. Western Blot results yielded 2 different bands, suggesting there are two states of ALP. SBA Lectin Blot (Figure 1B) analysis confirm that the second states were glycosylated. Glycosylation rates were estimated by calculating the light intensities coming from each band within the lanes in Western Blot (Figure 1C). According to the results, ALP-DQNAT was estimated to be the highest glycosylated construct with 45% glycosylation. Moreover, repeating the recognition motif did not increase the glycosylation rate further. On the contrary, a slight decrease was observed.

**Figure 1:**
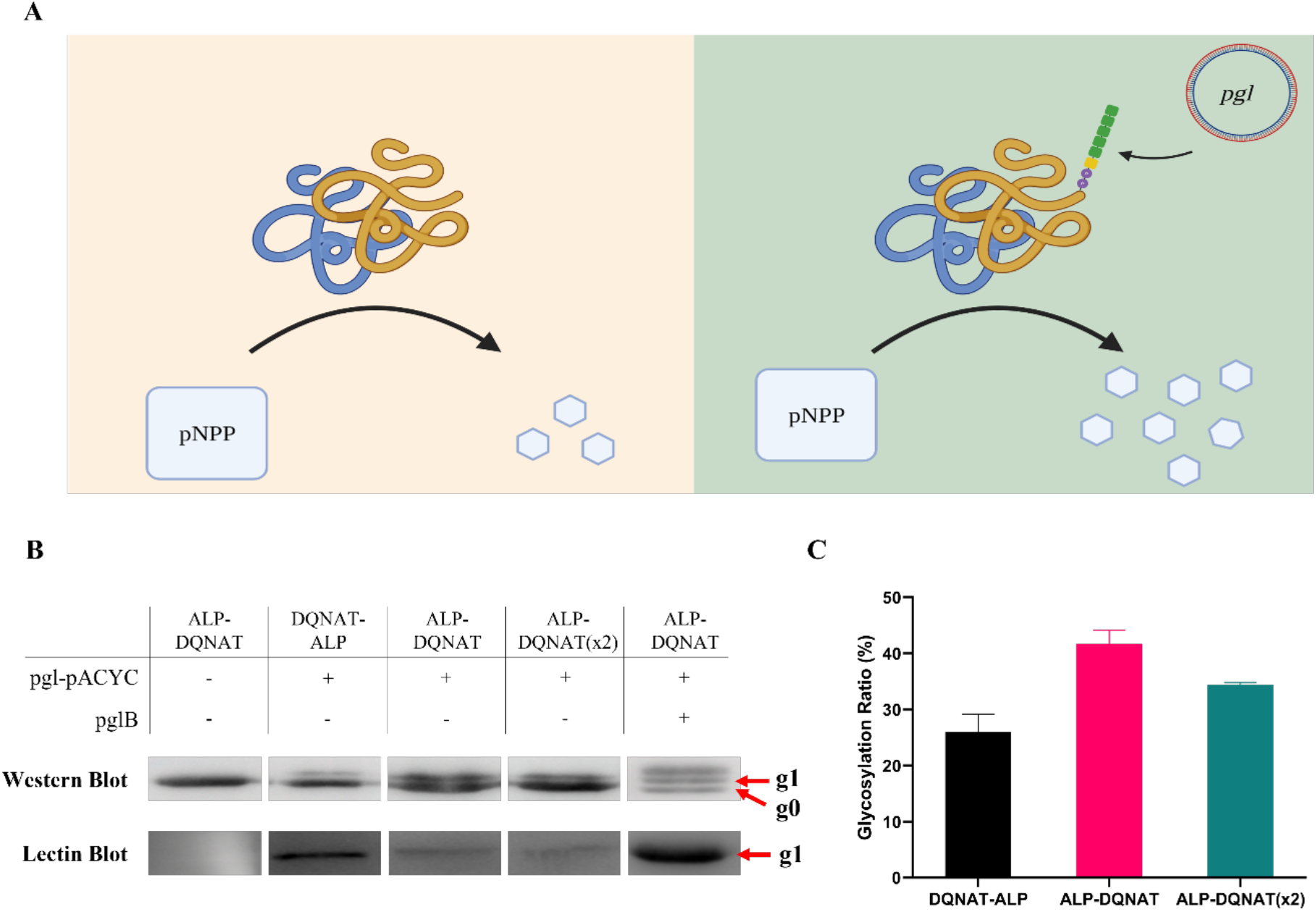
(A) Illustration of the effect of recombinant glycosylation on ALP enzyme. Created with BioRender.com (B) Western and lectin blot results of the designed contructs. Non-glycosylated ALP is denoted as “g0” and glycosylated ALP is denoted as “g1”. (C) Calculated glycosylation rate of the constructs based on the western blot results.

### Improving glycosylation efficiency in the cell

*N-linked glycosylation* machinery adds glycan groups onto target proteins in the periplasm of the cell. To eliminate ALP in the cytoplasm which is certainly unglycosylated, periplasmic protein extraction by osmotic shock has been performed. Western Blot analysis indicated that glycosylation rate does not change significantly (Figure S1), suggesting that most of ALP that is produced successfully translocates to periplasm. Therefore, to increase the glycosylation efficiency in the periplasm, we overexpressed PglB enzyme in the cell. Western Blot results yielded an extra band (Figure 1B, Lane 5) suggesting three states. Lectin Blot results contradicted and indicates there is only one glycosylated state. Therefore, the extra band most likely does not correspond to *N-glycosylated* ALP. However, after purification of the protein with Histag column, the extra band was eliminated during the purification process. The elution was checked with Western Blot and Lectin Blot to confirm the glycosylation and with SDS-PAGE for purity (Figure S2). After protein is purified, approximately 52% glycosylated batch of ALP was obtained and will be referred as “Glycosylated ALP-DQNAT” from now on.

### Assessing the enzymatic activity of glycosylated ALP

After achieving successful glycosylation of ALP and increasing the glycosylation rate, we investigated the effect of recombinant glycosylation on enzyme’s behavior. Para-Nitrophenylphosphate (pNPP) is the substrate of alkaline phosphatase enzyme and after it is cleaved by ALP, it forms a yellow-colored product. The alkaline phosphatase activity can be assessed by the rate of the color change in a transparent medium.(Sayler et al., 1979). We performed enzyme activity assay to assess the effect of the glycosylation at optimal conditions. The results showed that glycosylated ALP performed 1.5-fold better than the native enzyme (Figure 2A). We also performed Michealis-Menten analysis to evaluate the differences (Figure 2B). The analysis showed that ν_max_ is 1.0376 μM/min and 2.088 μM/min for ALP-DQNAT and glycosylated ALP-DQNAT, respectively. K_m_ value was also increased from 180.2 μM to 290 μM, upon glycosylation. Furthermore, to show the substrate conversion into product over time and calculate the average velocity, product formation versus time graph was drawn (Figure 2C). The average velocities were calculated as 21 nM s^-1^ and 34 nM s^-1^, respectively.

**Figure 2:**
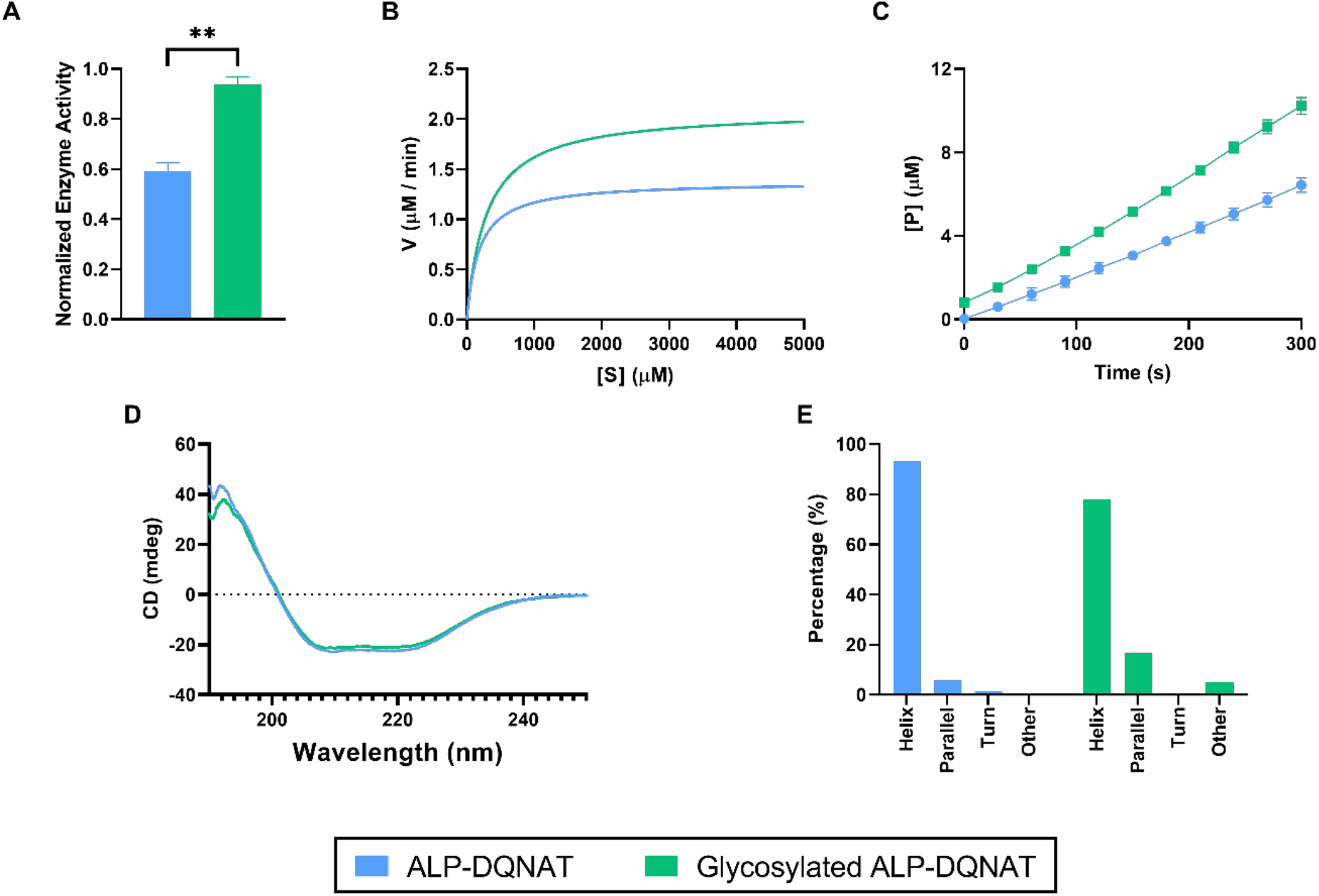
Analysis of ALP-DQNAT and Glycosylated ALP-DQNAT at optimal conditions. (A) Enzyme activity of ALP-DQNAT and glycosylated ALP-DQNAT. (B) Michealis-Menten analysis of ALP-DQNAT and glycosylated ALP-DQNAT. (C) The plot of substrate conversion performed by ALP-DQNAT and glycosylated ALP-DQNAT over time. (D) CD spectra results of ALP-DQNAT and glycosylated ALP-DQNAT. (E) Secondary structure predictions of ALP-DQNAT and glycosylated ALP-DQNAT using CD data.

To understand the changes in the activity of the enzyme, changes in the secondary structures upon glycosylation was also analyzed. Circular Dichroism (CD) is a widely utilized method to study secondary structures of proteins (Greenfield, 2006). To determine the secondary structures, a wavelength range between 190 nm and 250 nm was utilized. Results showed that *N-linked glycosylation* did not affect the CD spectra significantly and ALP (Figure 2D). Similar results have been reported previously (Sahin Kehribar et al., 2021; Wu et al., 2017). To predict the secondary structures BestSel online tool was utilized(Micsonai et al., 2018). Results indicate that no significant change was observed, and ALP preserved its alpha-helical structure (Figure 2E).

### *N-linked glycosylation* increases the stability of ALP enzyme

To test the effect of glycosylation on the enzyme activity at elevated temperatures, both enzymes were incubated at 55°C, 75°C and 95°C for varying incubation times (15 min, 30 min, 60 min and 120 min) (Figure 3A). At 55°C, activity of both enzymes were decreased compared to untreated groups in Figure 2A. However, glycosylated ALP-DQNAT preserved 55% of its activity, whereas ALP-DQNAT showed 29% of its activity at optimal conditions. At 75°C, ALP-DQNAT enzyme become susceptible to incubation time as well and continue losing its activity as incubation time increases. Interestingly, glycosylated ALP-DQNAT preserved its activity in increasing incubation times. Therefore, the fold change between ALP-DQNAT and glycosylated ALP-DQNAT was 2-fold when treated for 15 minutes, it rose to 6.1-fold for 120 minutes-treatment. At 95°C, both enzymes cease to work after 30 minutes of incubation. Overall, the results indicated that glycosylated ALP-DQNAT performed better at all conditions. We also investigated the effect of elevated temperatures on the secondary structure of both enzymes. Both enzymes were treated with elevated temperatures (55°C, 75°C, 95°C) for 2 hours. Measurements were taken immediately after treatment. CD results were used as input for BestSel tool and the secondary structure predictions were shown in Figure 3B. According to the results, 55°C and 95°C did not cause a significant change in the secondary structure. However, at 75°C, parallel structure of glycosylated ALP was diminished and it was compansated by helix and other secondary structures. This change in the secondary structure may explain the reason why ALP-DQNAT activity was weakened over time, whereas glcosylated ALP-DQNAT protected its activity.

**Figure 3:**
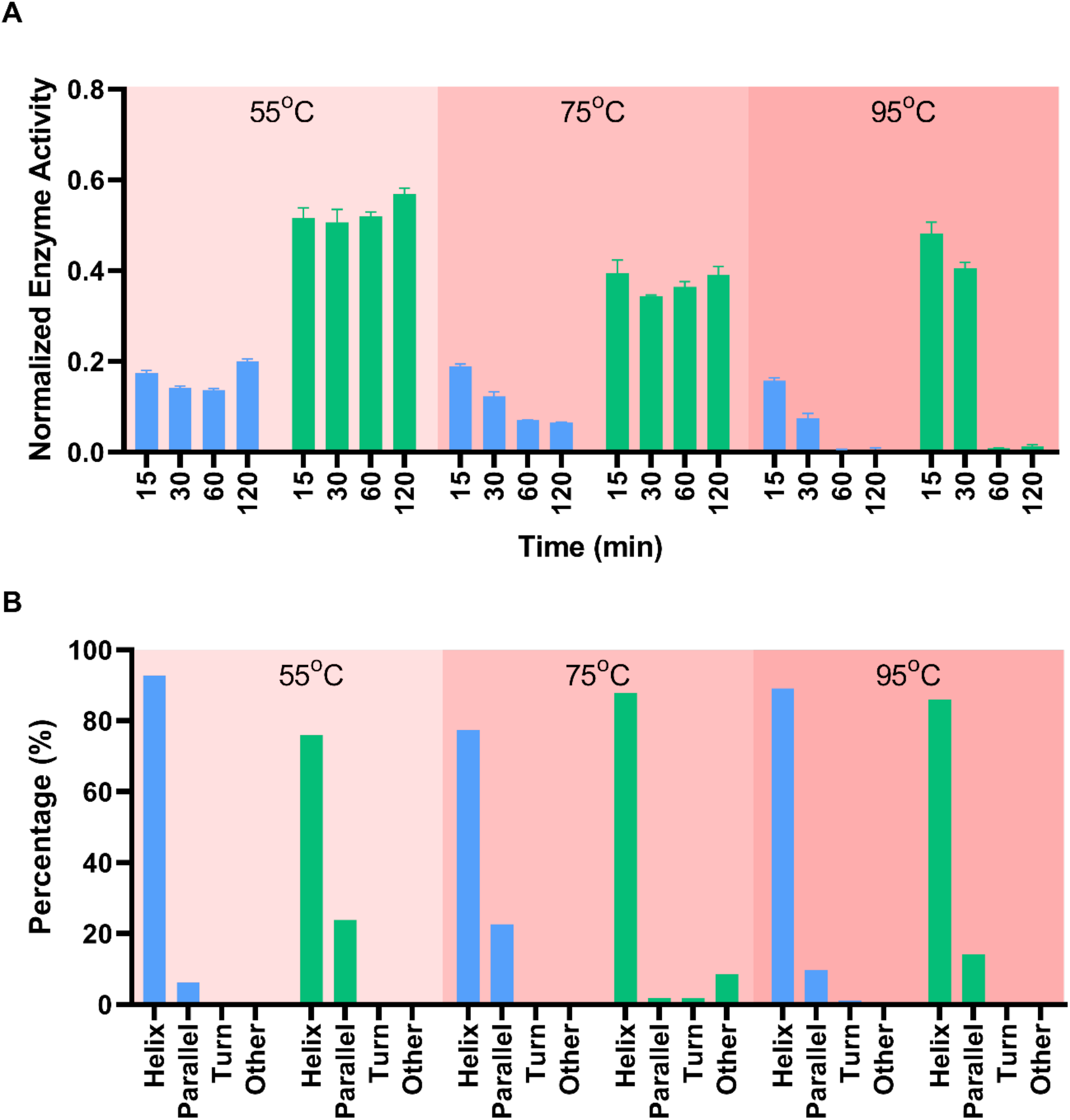
Glycosylation leads to better performing ALP. (A) Enzyme activity results of ALP-DQNAT and glycosylated ALP-DQNAT incubated at elevated temperatures. (B) Secondary structure predictions of ALP-DQNAT and glycosylated ALP-DQNAT.

### Glycosylated ALP-DQNAT is more active at suboptimal pH conditions and more alkaline pH

pH is one of the most significant effectors of enzyme activity. Therefore, pH conditions should be well arranged for enzymes to work efficiently. *E. coli* alkaline phosphatase enzyme has optimum pH of 8.0 (Garen and Levinthal, 1960). To examine how glycosylation affects the working pH range of the enzyme, we performed enzyme acitivity assays at different pH conditions, ranging from pH 5.0 to 13.0 (Figure 4A). At pH 5.0, 6.0, 12.0 and 13.0, no enzyme activity was detected. At other tested pH conditions, glycosylated ALP-DQNAT activity was higher, compared to ALP-DQNAT. Furthermore, it can be said that, upon glycosylation, the optimum pH for the enzyme has shifted to more alkaline conditions, as the highest activity was seen at pH 10.0. We also studied secondary structures at pH conditions where enzyme activity was observed (Figure 4B). However, no significant difference that may account for enzyme activity results was observed.

**Figure 4:**
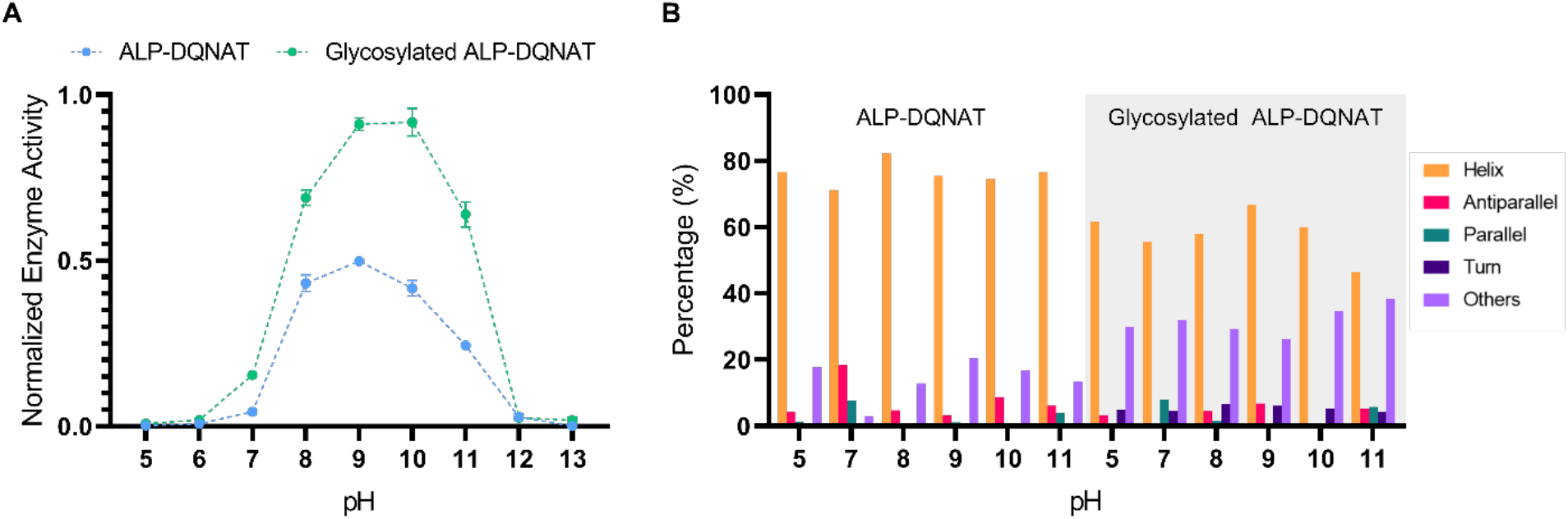
Working pH range screening for ALP-DQNAT and glycosylated ALP-DQNAT. (A) Glycosylated ALP-DQNAT performs better for all pH conditions tested. (B) Secondary structure predictions of ALP-DQNAT and glycosylated ALP-DQNAT

### Glycosylation protects ALP against proteolytic cleavage

Another important characteristic of proteins is stability against proteolytic cleavage. Proteases are enzymes that degrades proteins into smaller peptides. We utilized a serine protease, Proteinase K, to assess the stability of enzymes against proteases. Proteinase K has also optimum pH of 8.0, therefore, it is convenient to use in the same medium.

We first checked proteolytic cleavage on SDS gel (Figure 5A). SDS gel confirms that both enzymes are subjected to degradation. It was observed that both proteins were not able to protect their full size. We also investigated the effect of incubation time with PrK (Figure 5B). It was seen that 60 minutes treatment is sufficient to observe the effects of PrK. Therefore, incubation time was selected as 60 minutes for concentration dependent analysis. Next, we applied increasing concentrations of PrK (Figure 5C). It was observed that, the activity fold change increases as PrK concentration increases. At 10 μg/mL, the fold change increases to 24-fold, where ALP activity is very limited. Therefore, it can be stated that glycosylation contributes to stability of ALP against proteases.

**Figure 5:**
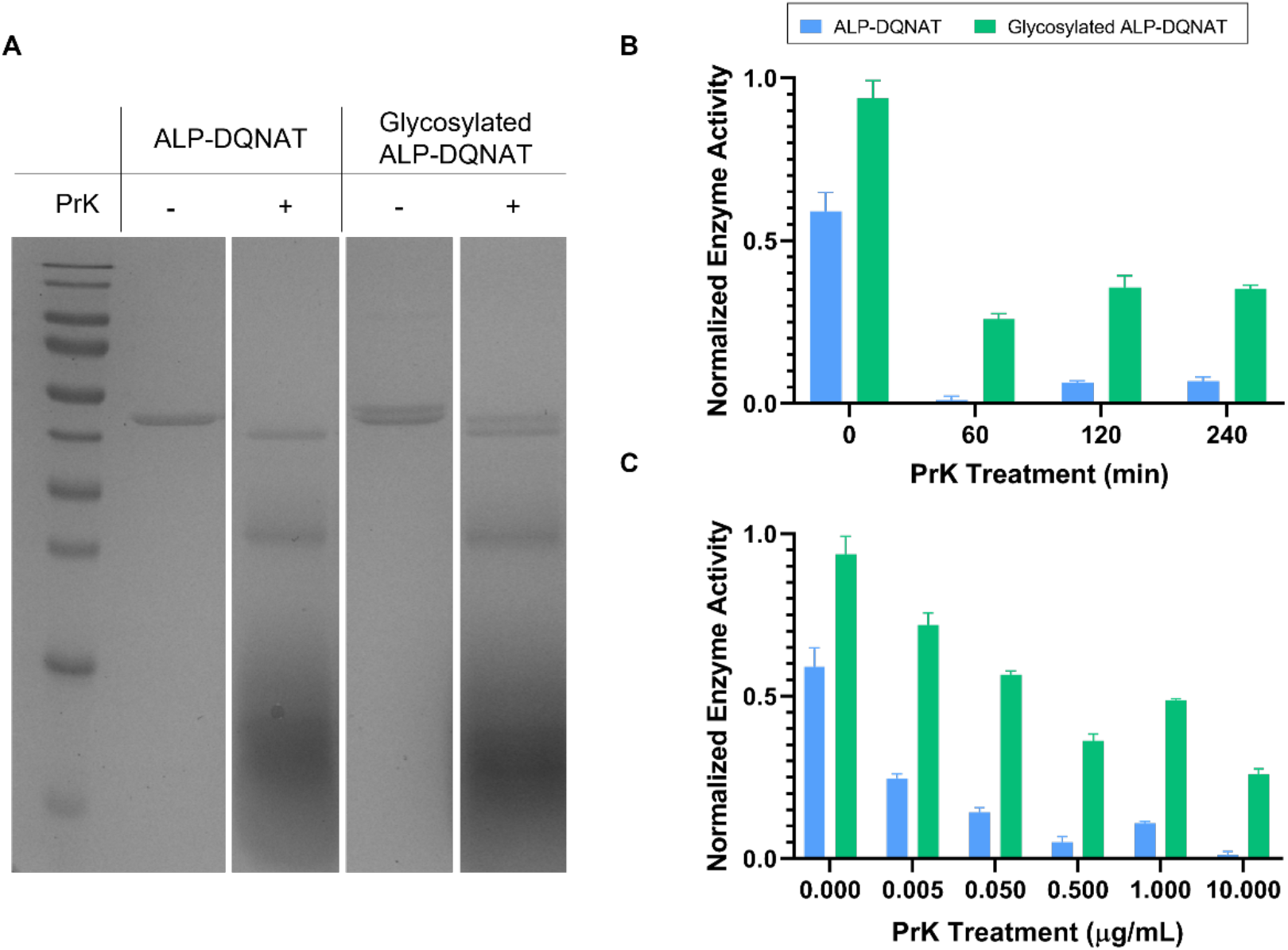
ALP-DQNAT and glycosylated ALP-DQNAT is assessed in terms of their stability against proteolytic cleavage. (A) SDS-PAGE analysis was performed to observe proteolytic cleavage performed by PrK. (B) ALP-DQNAT and Glycosylated ALP-DQNAT were treated with PrK for varying incubation times. (C) Increasing PrK was applied to both enzymes to investigated the alterations in enzymatic activity.

### Enzyme activity varies under denaturing conditions

We investigated the enzymatic behaviours of both ALP-DQNAT and glycsoylated ALP-DQNAT under denaturant conditions as well. We utilized two common denaturing agents, guanidine hydrochloride and urea. We first started examining the effects of urea on enzyme activity (Figure 6A). We observed that in urea, enzyme activity of ALP-DQNAT is higher at all conditions tested. We also wanted to examine the effect of guanidine on enzyme acitivities to understand whether the results we observed are specific to urea. Addition of guanidine showed that glycosylated ALP-DQNAT performed better at all concentrations applied (Figure 6B). At moderate concentrations (50 mM and 100 mM), enzyme activities were enhanced. At high concentrations, ALP-DQNAT lost its enzymatic activity completely whereas, glycosylated ALP-DQNAT retained meaningful activity levels. Therefore, glycosylation enhanced the stability of ALP-DQNAT against guanidine.

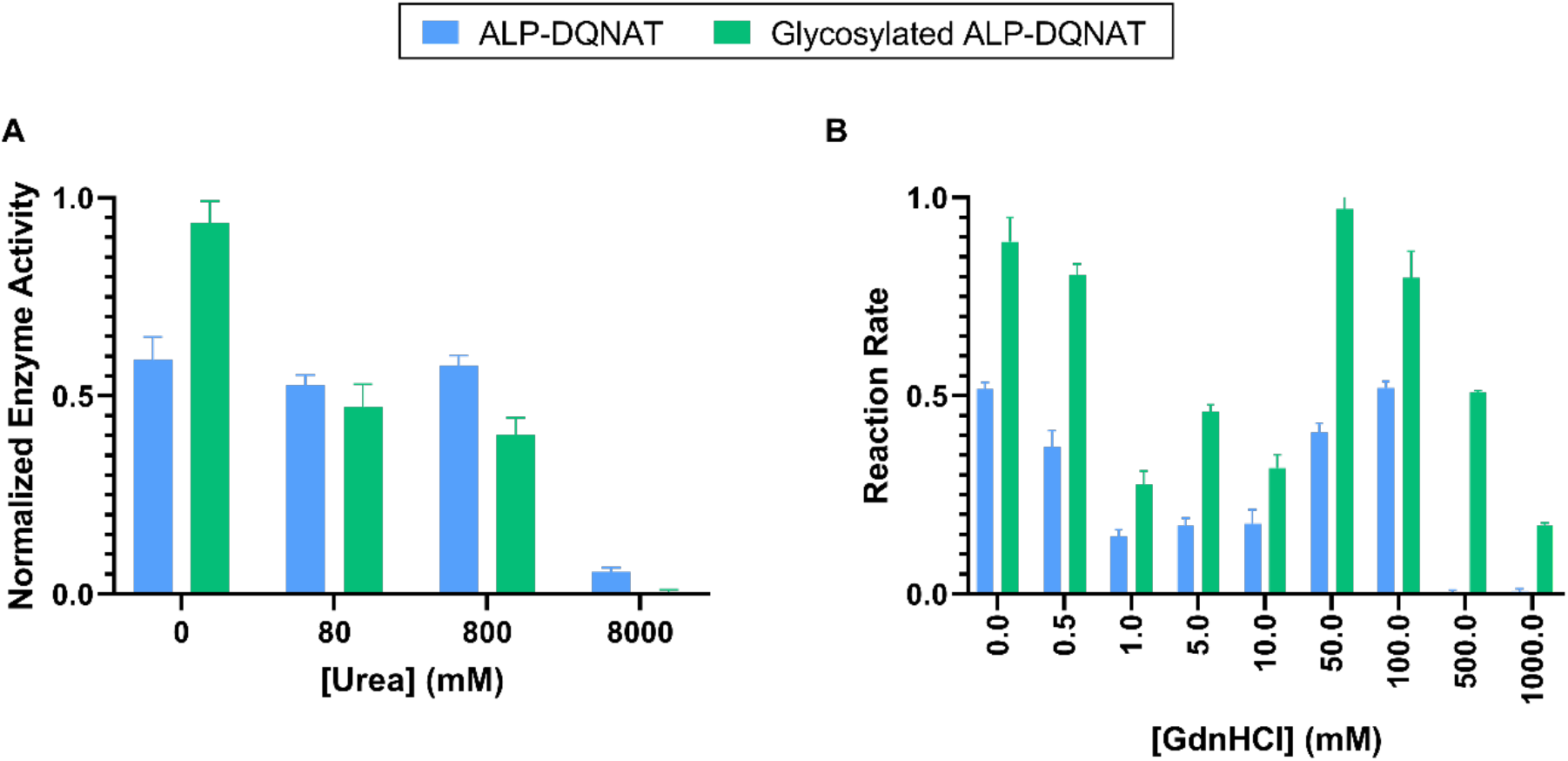

## Discussion

In this work, we exploited the effect of N-linked glycosylation on enzyme’s behaviour, which has not been investigated before. To accomplish this, we designed different constructs that encodes ALP enzyme with the glycosylation tag at different locations. Since the success of glycosylation depends on the accessibility of glycosylation tag by PglB oligosaccharyltransferase, each construct led to different glycosylation rate (Silverman and Imperiali, 2016). ALP enzyme was successfully glycosylated, highest glycosylated construct was selected, and overexpressed in *E. coli*. To increase this glycosylation rate further, PglB enzyme was overexpressed. Overexpression of PglB resulted in an “extra” band in western blot (Figure 1b). However, the “extra” band was not confirmed by lectin blot, making it miscellaneous. The simplest explanation could be that ALP-DQNAT is di-glycosylated at a different site. To explore this possibility, we used an *in silico* tool GlycoPP to uncover putative N-linked glycosylated sites (Chauhan et al., 2012). GlycoPP returned that three more sites is potentially N-linked glycosylated (Figure S3). Similar results were reported in the literature before (Ollis et al., 2015). Another possibility is that, another or inmature glycan is attached to ALP-DQNAT. It has been previously reported by many studies that PglB has relaxed specificity (Feldman et al., 2005; Wacker et al., 2006). Although, it is known that *C. jejuni* utilizes only one type of oligosaccharide, inmature glycans could be attached to one or more of the potential glycosylated sites. In this way, lectin blot could not detect the inmature glycans, therefore, a slower migrating band was observed. In the same manner, O antigens could be attached to ALP-DQNAT as well and they also could escape lectin blot, resulting in the same situation, since the strain utilized already synthesize O antigens (Feldman et al., 2005).

We demonstrated that glycosylated ALP shows better performance, compared to native enzyme, in terms of catalytic efficiency. There are different approaches to compare enzymes that catalyzes the same reaction. Michealis constant is vastly utilized to assess the performance of enzyme’s over the last century (Kim and Tyson, 2020). While it is useful to evaluate enzymes in their natural environments, it does not take into account industrial processes such as high substrate value, non-physiological pH and temperature values (Carrillo et al., 2010). Furthermore, with the advances in genetic engineering such as directed evolution, comparing enzyme variants acting on the same substrate become tricky utilizing *k*_*cat*_/*K*_*M*_ (Eisenthal et al., 2007). Therefore, new evaluation methods and parameters are required. Since the important parameter is the completion of the reaction in most of the industrial processses, tracking of product formation and comparing enzymes accordingly would be more suited (Fox and Clay, 2009).

We also showed that glycosylation makes ALP-DQNAT more durable. There has been no evidence that *C. jejuni* N-linked glycosylation contributes to thermostability of the enzymes. However, it is well known that glycosylation in general may alter the stability of proteins and it was also shown at elevated temperatures (Eriksen et al., 1998; Khan et al., 2003; Sola and Griebenow, 2009). In line with the literature, our results clearly indicates that glycosylation improves thermostability of ALP. Aside from thermostability, N-linked glycosylation was stated to contribute to stability against proteases(Dotsenko et al., 2016; Kightlinger et al., 2020). Glycosylation provides this protection by shielding protein regions from proteases (Sola and Griebenow, 2009).As in previous reports, we found that glycosylated ALP-DQNAT outperforms ALP-DQNAT. Furthermore, we examined enzyme activity at different pH conditions. We observed a slight shift to more alkaline pH conditions upon glycosylation. This is in good agreement with other studies that examines the effect of glycosylation (Dotsenko et al., 2016; Guo et al., 2008).

We also tested both enzymes under denaturing conditions, urea and guanidine hydrochloride. Surprisingly, addition of urea reversed the fashion followed in the enzymatic activities and diminished the activity of glycosylated ALP more, compared to unglycosylated ALP. Later we found out that, N-glycans react with high concentrations of urea and lead to formation of artefact (Omtvedt et al., 2004). Therefore, this may partially explain the decrease in enzyme activities. Then, we also examined enzyme acitivities in Gdn-HCl. Enzymes performed as expected with the addition of Gdn-HCl. It has been clearly seen that glycosylation ALP-DQNAT performed better at all conditions and continue working where ALP-DQNAT activity was lost. A surprising result was observed at moderate concentrations. Moderate concentrations of Gdn-HCl lead to an unexpected stimulation in both enzyme’s activities.

This effect of guanidine on ALP has been previously found and discussed and therefore, results are in agremeent with the literature (Rao and Nagaraj, 1991).

Overall, presented results above showed that glycosylation of ALP-DQNAT added a significant value to enzyme’s characteristics and it is a promising strategy to improve enzymes to be utilized in industrial processes.

## Significance

Recombinant enzymes are becoming more important in many fields, including medicine and industrial processes. Enzymes usually work fine in their natural environment, however, they do not meet the expectations in the realm of bioprocess conditions, such as non-physiological pH, temperature and exposure to non-natural solvents. Therefore, they need to undergo an optimization process to be suited for their specific applications. Here, we propose a useful method to tailor enzymes to make them better suited for industrial processes. Utilizing N-linked glycosylation, we showed that glycosylated ALP is better performing enzyme. We increased its thermostability, which is one of the major parameters, leading 6.1-fold increase in enzyme activity. Another parameter, pH, was also examined and we demonstrated that glycosylated ALP activity is more stable, compared to unglycosylated counterpart. We also investigated the stability against proteases and our results clearly showed significant improvements in enzyme acitivity in the presence of proteases. Lastly, enzymes were tested in denaturing conditions and we showed that glycosylated ALP is superior to unglycosylated ALP. With our results, we anticipate that N-linked glycosylation of enzymes allow a rapid optimization of enzymes for industrial processess and offers improved features of enzymes in terms of temperature, pH, proteolytic stability and denaturing conditions.

## Notes

### Competing Interest Statement

The authors have declared no competing interest.

